# A polycomb-independent role of EZH2 in TGFβ1-damaged epithelium triggers a fibrotic cascade with mesenchymal cells

**DOI:** 10.1101/2020.07.29.225300

**Authors:** Huy Quang Le, Matthew Alexander Hill, Ines Kollak, Wioletta Skronska-Wasek, Victoria Schroeder, Johannes Wirth, Eva Schruf, Karsten Quast, Franziska Elena Herrmann, Matthew James Thomas, James Peter Garnett

**Author notes:** Correspondence: H.Q.L.

## Abstract

To restore organ homeostasis, a myriad of cell types need to activate rapid and transient programs to adjust cell fate decisions and elicit a collective behaviour. Characterisation of such programs are imperative to elucidate an organ’s regenerative capacity and its aberrant repair in disease. By modelling epithelial-mesenchymal crosstalk, we provide direct evidence for transforming growth factor β1 (TGFβ1)-damaged epithelium initiating a bi-directional fibrotic cascade with the mesenchyme. Strikingly, TGFβ1-damaged epithelia facilitates the release of Enhancer of Zester Homolog 2 (EZH2) from Polycomb Repressive Complex 2 (PRC2) to establish a novel fibrotic transcriptional complex of EZH2, RNA-polymerase II (POL2) and nuclear actin. Perturbing this complex by disrupting epithelial EZH2 or actomyosin remodelling abrogates the fibrotic crosstalk. The liberation of EZH2 from PRC2 is accompanied by an EZH2-EZH1 switch to preserve global H3K27me3 occupancy. Our results reveal an important non-canonical function of EZH2, paving the way for therapeutic interventions in fibrotic disease.

## Main

How organisms activate repair and regeneration programs upon injury to restore physiological conditions is a fundamental question in biology. This process arises from communication between several cell types (notably epithelial cells, endothelial cells, immune cells, mesenchymal cells and neurons) to coordinate changes in their gene expression and behaviour that lead to tissue remodelling^1^. Cells sense and transmit these signals through receptors, cell-cell and cell-extracellular matrix (ECM) interactions that couple extrinsic signals through the actomyosin cytoskeleton to the nucleus and eventually the chromatin. Any dysregulation may result in the transition to disease, e.g. cancer or fibrosis^2,3^.

Idiopathic Pulmonary Fibrosis (IPF) is a chronic respiratory disease characterized by progressive fibrotic lung remodelling and respiratory failure. The disease is ultimately fatal, despite the emergence of current therapeutics^4^. IPF initiation and progression have been linked to damage and remodelling of the respiratory epithelium by evidence derived from genetic screening, animal models of fibrosis and pathological analyses of patient lungs^1,5^. Although these studies suggest that epithelial secretions and their crosstalk with fibroblasts are key drivers of IPF disease pathology, the exact underlying mechanisms remain to be established^6^.

TGFβ1 is a predominant effector in most, if not all, forms of fibrosis. Several studies have uncovered an intricate crosstalk between SMADs, the central player of TGFβ signalling, and a myriad of epigenetic regulators to fine-tune the transcriptional machinery^7,8^. PRC2 is a major epigenetic repressive regulator consisting of EZH2, SUZ12 and EED. PRC2 catalyses mono-, di- and tri-methylation of Histone 3 on Lysine 27 (H3K27me2/3) through its methytranferase EZH2 subunit resulting in gene silencing and chromatin compaction^9^. Importantly, EZH2 overexpression has been detected in IPF and cancer showing a positive correlation with their progression^10-12^. Intriguingly, a non-canonical role of EZH2, marked by phosphorylation on threonine 311 (T311) and/or T487, in which it acts as a transcriptional activator via a PRC2- and methylation-independent manner, has been identified in several cancers^13,14^. Nonetheless, the molecular principles dictating the role of EZH2 as either a transcriptional silencer or activator is far from understood. Here we describe for the first time a PRC2-independent role of EZH2 as a transcriptional co-activator during TGFβ1-driven expression of pro-fibrotic genes in human lung epithelial cells. This requires liberation of EZH2 from the PRC2 complex, followed by interactions between EZH2, POL2 and actin. Simultaneously, EZH1 is recruited and forms an EZH1-PRC2 complex to maintain the silencing of non-target genes. These changes trigger a pro-fibrotic crosstalk with mesenchymal cells, leading to ECM remodelling and further damage of the epithelium. Characterisation of this pathologic network provides an opportunity for novel therapeutic intervention through inhibition of non-canonical EZH2.

To mimic epithelial-mesenchymal crosstalk in the human lung during fibrogenesis, we utilized a co-culture system wherein human lung alveolar epithelial cells A549 (EPCs)^15^ were apically treated with TGFβ1 on Transwell inserts, and cultured with either primary normal human lung fibroblasts (NHLFs) or IPF-derived lung fibroblasts (IPF-LFs) at the bottom of the culture well for 72 h (Fig. 1a). Apical TGFβ1 treatment of EPCs within the co-culture system led to loss of E-cadherin (E-cad), a marker for epithelial integrity (Fig. 1b,c). This coincided with enhanced alpha-smooth muscle actin (α-SMA) and other pro-fibrotic/ECM proteins levels including fibronectin (FN), type I collagen (COL-1), matrix metalloproteinase-7 (MMP7) and Monocyte Chemotactic Protein 1 (MCP1) in the mesenchymal compartment (Fig. 1d and Supplementary Fig. 1a-d). Importantly, these pro-fibrotic responses to TGFβ1 were only observed in the presence of an epithelial layer, while no elevation of TGFβ1 could be detected in the mesenchymal compartment after 72 h (Supplementary Fig. 1e,f), indicating that signals from damaged epithelium beyond secreted TGFβ1 drive FMT (Fibroblast-to-Myofibroblast Transition)/pro-fibrotic effects in our co-culture model. Interestingly, this effect was even more pronounced in IPF-derived fibroblasts (Fig. 1e).

**Figure 1.**
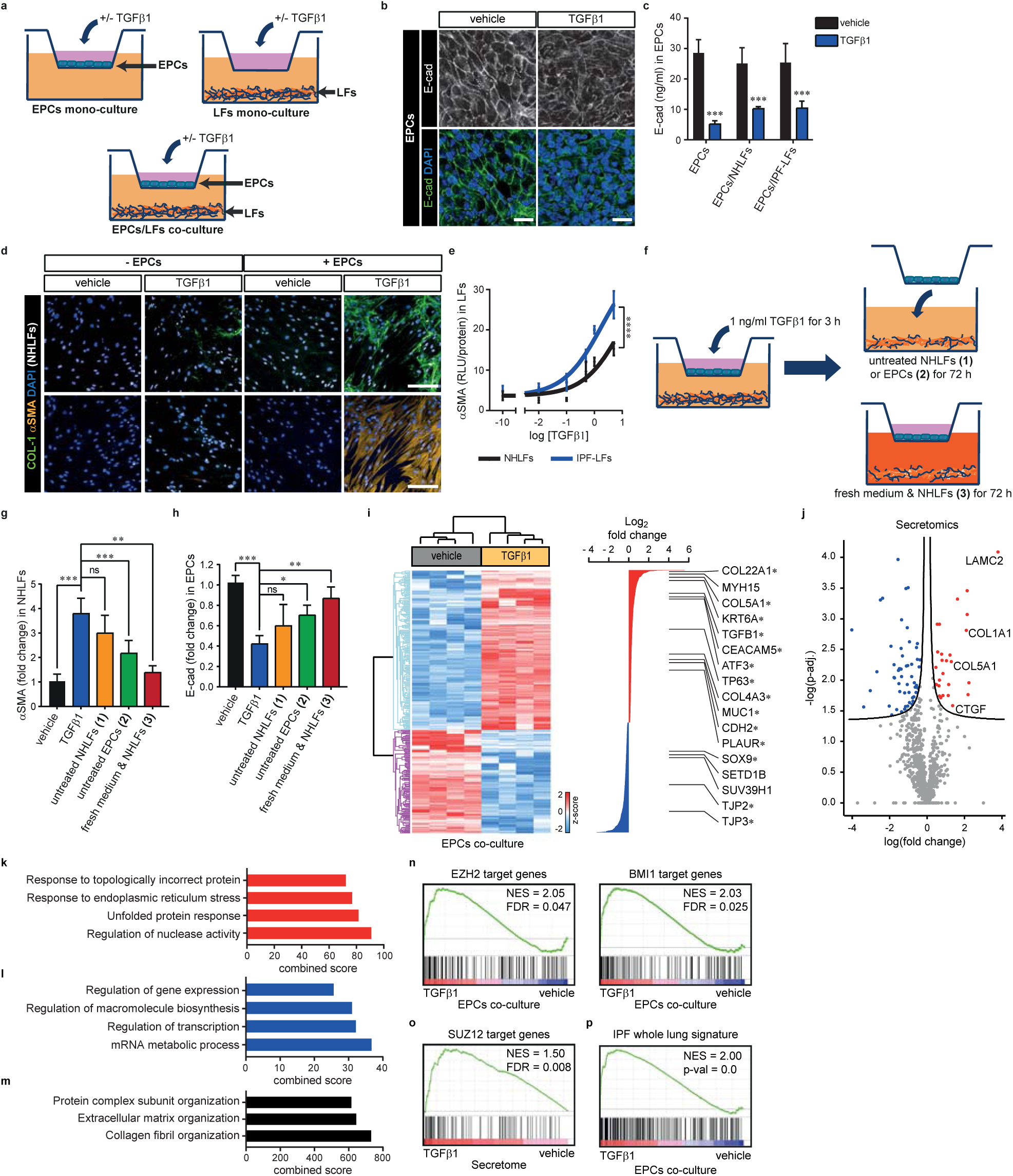
Polycomb mediates TGFJ31-induced epithelial secretions triggers bi-directional fibrotic cascade with mesenchymal cells. **(a)** EPCs were subjected to apical TGFβ1 stimulation for 72 h in the co-culture system. **(b)** Representative E-cadherin (E-cad) and DAPI immunofluorescence images of EPCs treated apically with TGFβ1 or vehicle control after 72 h in the co-culture system (scale bars 25μm). **(c)** ELISA analysis shows decreased E-cad levels in TGFβ1-treated EPCs after 72 h in the mono- and co-culture systems (n = 6 independent experiments, mean + s.d., ***p < 0.001, ANOVA/Tukey’s). **(d)** Representative COL-1, αSMA and DAPI staining of NHLFs from the mono- and co-culture systems after 72 h apical treatment with TGFβ1. Note the increased COL-1 levels, further enhanced by the addition of EPCs and an increase in αSMA levels only in the presence of EPCs (scale bars 200μm). **(e)** ELISA analysis shows increased αSMA levels in a TGFβ1 dose-dependent manner from NHLFs and IPF-LFs in the co-culture system. Note a stronger αSMA response in IPF-LFs (mean + s.d., n = 3 independent experiments from 6 NHLFs and 6 IPF-LFs donors, ****p < 0.0001, Nonlinear Regression). **(f)** Schematic workflow of the substitution system in which EPCs were subjected to TGFβ1 stimulation for 3 h in co-culture with NHLFs, after which (1) NHLFs or (2) EPCs were replaced with untreated counterparts or (3) both fresh medium and untreated NHLFs for a further 72 h. **(g, h)** 72 h after substitutions in the co-culture, αSMA and E-cad levels of NHLFs and EPCs, respectively, were measured with ELISA and normalised to the control. Note replacement of injured EPCs with untreated EPCs or replacement of media and NHLFs attenuates αSMA expression and increases E-cad levels (mean + s.d., ***p < 0.001, **p < 0.01, *p < 0.05, n = 3 independent experiments with 6 NHLFs donors, ANOVA/Dunnett’s). **(i)** Hierarchical clustering of differentially regulated transcripts from RNA-seq between vehicle and TGFβ1-treated EPCs in the co-culture (p-adj < 0.05). Significantly regulated pro-fibrotic genes (marked by asterisk) are listed on the right side. **(j)** Volcano plot representing logarithmic ratio of differentially secreted proteins from proteomics analysis of co-culture medium upon apical TGFβ1 stimulation (p-adj < 0.05), with examples of pro-fibrotic proteins. **(k-m)** Gene Ontology (GO) analysis of differentially expressed genes (p-adj < 0.05) that are enriched in TGFβ1-treated (k), vehicle treated (l) EPCs, and differentially secreted proteins (m). **(n, o)** GSEA shows enrichment of genes that are known targets of Polycomb complexes (defined by SUZ12, BMI1 and EZH2) upon TGFβ1 treatment in the co-culture system. **(p)** GSEA shows enrichment of an IPF whole lung signature in TGFβ1-treated EPCs in the co-culture system.

As fibroblasts have been shown to shape epithelial homeostasis^16^, we next assessed whether activated fibroblasts in turn influence EPCs. To test this, we injured EPCs with TGFβ1 for 3h in co-culture with NHLFs, followed by substitution with either (1) untreated NHLFs, (2) untreated EPCs, or (3) fresh medium & NHLFs for a further 72 h (Fig. 1f). We observed an induction of α-SMA levels in substituted NHLFs (1), consistent with epithelial-derived pro-fibrotic secretions driving FMT. Accordingly, replacement with healthy EPCs (2) or combination of fresh medium & NHLFs (3) decreased α-SMA expression in NHLFs (Fig. 1G). Furthermore, the untreated EPCs substituted into the NHLFs co-culture, exhibited a significant decrease in E-cad levels (2), indicating that activated fibroblasts damage the epithelium in return (Fig. 1h). Collectively, these experiments demonstrate a TGFβ1-initiated bi-directional fibrotic cascade in which injured EPCs trigger the activation of fibroblasts, which induce further epithelial injuries, pro-fibrotic mediator release and enhanced FMT.

We next sought to identify the molecular programs that mediate the bi-directional crosstalk. After 72h of co-culture upon apical TGFβ1 treatment, transcriptional profiles of the epithelial and mesenchymal cells were analysed using RNA sequencing (RNA-seq). We found 1732 significantly differentially expressed genes (DEGs) in EPCs, whereas 3778 genes were deregulated in NHLFs upon apical TGFβ1 treatment (*q-*val < 0.05, Fig. 1i, Supplementary Fig. 1g and Supplementary Table 1). Furthermore, secretory proteins were also characterised from the medium using mass-spectrometry-based secretome profiling. In response to apical TGFβ1-treated EPCs/NHLFs in the co-culture, 60 significantly deregulated proteins (*q-*val < 0.05, Fig. 1j and Supplementary Table 2) were detected and showed a positive correlation with RNA-seq data (Supplementary Fig. 1i,j). Intriguingly, among the most enriched gene ontology (GO) terms were genes/proteins involved in the development and progression of fibrotic disease, including regulators of mRNA metabolic processes, the unfolded protein response and ECM^17^ (Fig. 1k-m and Supplementary Fig. 3).

**Figure 2.**
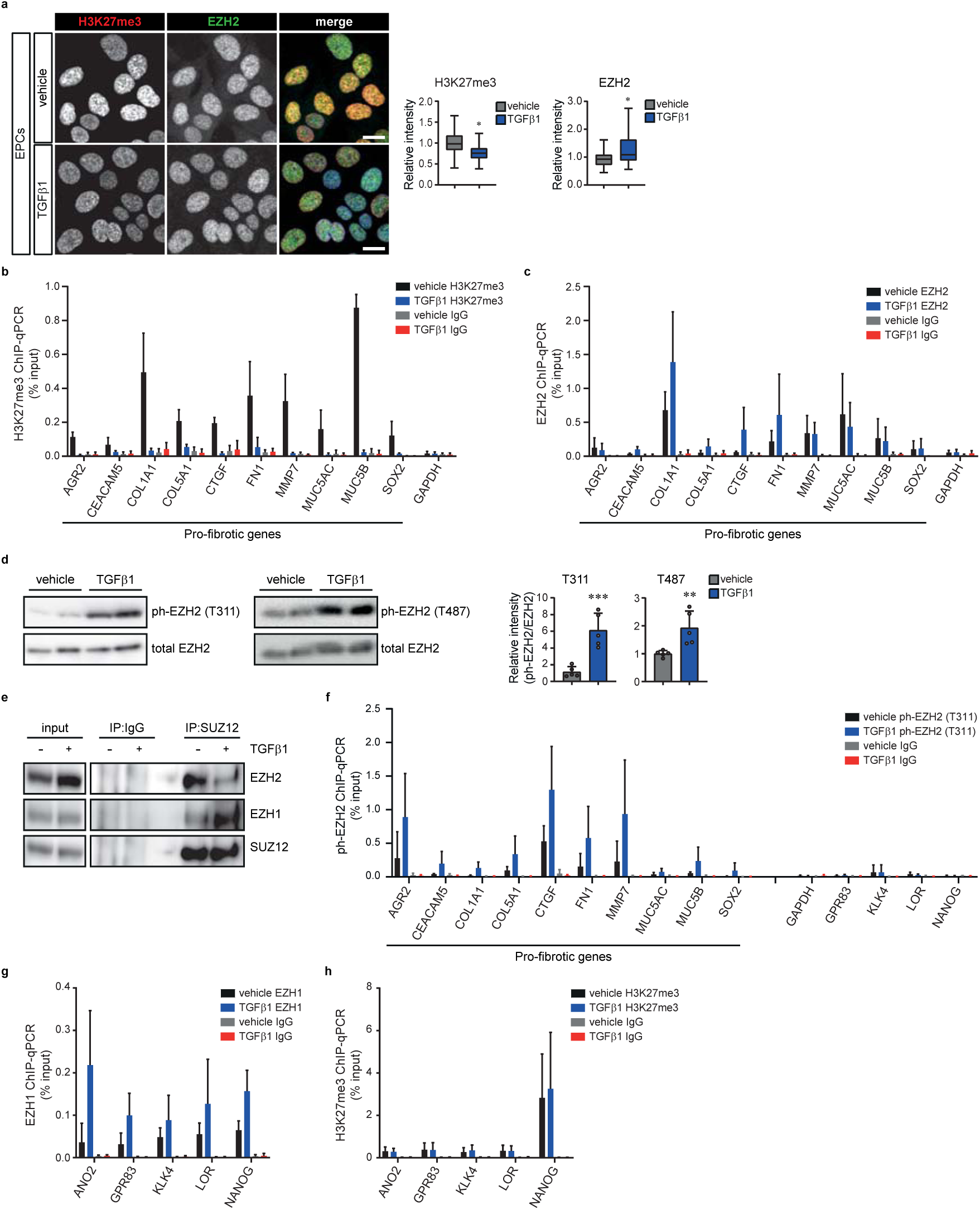
TGFβ1 induces an EZ-switch from EZH2 to EZH1-SUZ12 in EPCs of the co-culture system. **(a)** Representative H3K27me3 and EZH2 immunofluorescence images and box plots (minimum, first quartile, median, third quartile and maximum) showing decreased H3K27me3 levels but increased total EZH2 levels in EPCs from co-culture with LFs (n = 5 independent experiments with >50 cells per experiement, scale bars 50 m, *p = 0.0625, Wilcoxon test). **(b)** ChIP-qPCR shows decreased occupancy of H3K27me3 at promoters of pro-fibrotic genes in EPCs subjected to apical TGFβ1 for 72 h. Unspecific IgG was used as a negative control (mean + s.d., n = 3). **(c)** ChIP-qPCR shows increased EZH2 occupancy at promoters of profibrotic genes in EPCs subjected to TGFβ1 for 72 h (mean +s.d., n= 3). **(d)** Representative western blot analysis of ph-EZH2 and quantification shows increased ph-EZH2 levels at T311 and T487 in EPCs subjected to apical TGFβ1 for 72 h (n = 5 independent experiments, mean + s.d., **p = 0.0094, ***p = 0.0008, two-sided unpaired t-test). **(e)** Representative western blot analysis of SUZ12 immunoprecipitates shows co-precipitation of EZH1 and EZH2 in EPCs. TGFβ1 treatment leads to the EZ-switch from SUZ12-bound EZH2 to EZH1. Unspecific IgG binding was used as a negative control. A representative from 3 independent experiments is shown. **(f)** ChIP-qPCR shows increased ph-EZH2 occupancy at promoters of pro-fibrotic genes in EPCs subjected to TGFβ1 for 72 h. Note no changes in ph-EZH2 levels at promoters of non-target genes (mean + s.d., n = 3). **(g)** Increased EZH1 occupancy at promoters of non-fibrotic genes in EPCs subjected to apical TGFβ1 for 72 h (mean +s.d., n = 3). **(h)** No changes in H3K27me3 at promoters of non-fibrotic genes in EPCs subjected to apical TGFβ1 for 72 h (mean + s.d., n = 3).

**Figure 3.**
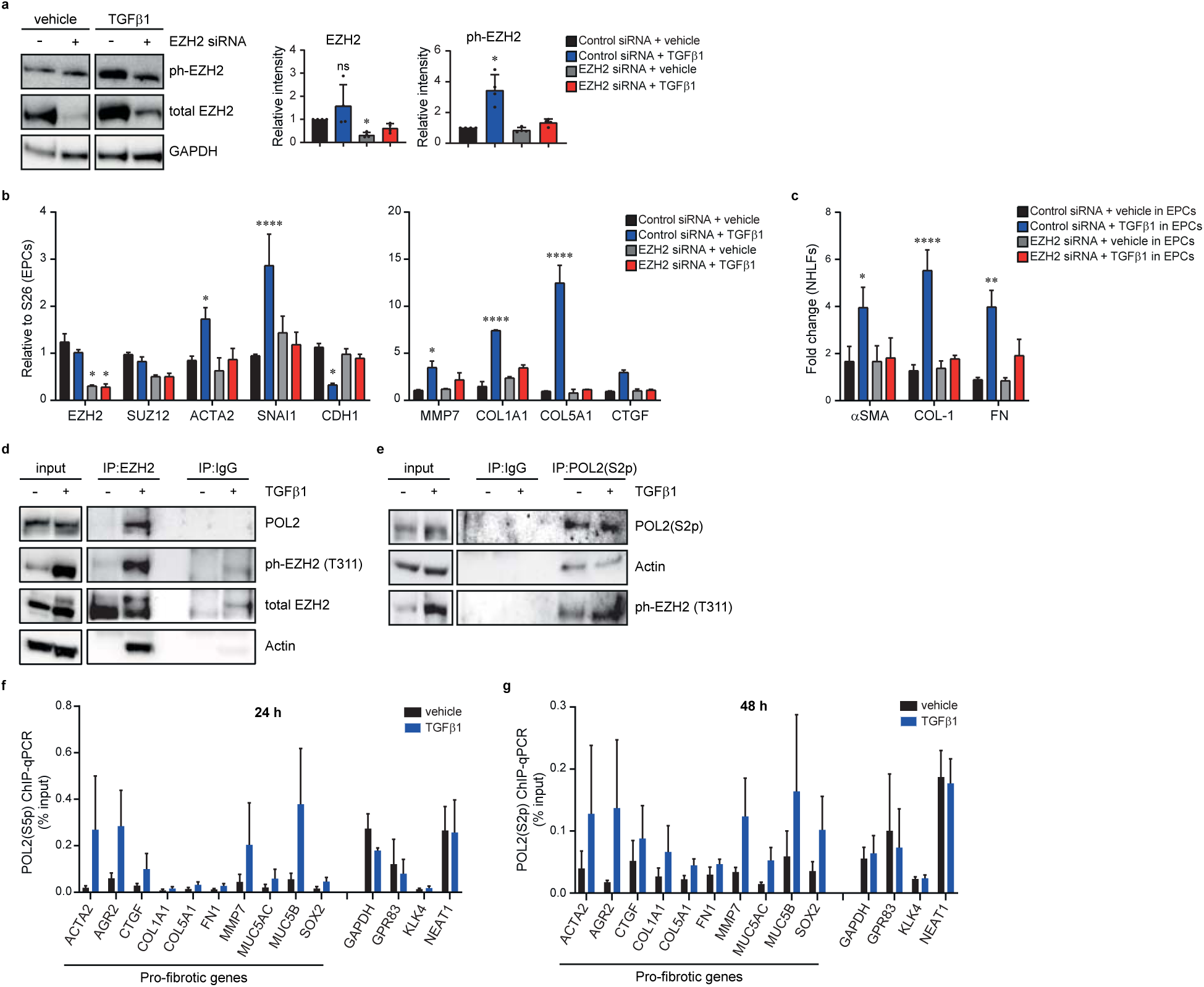
TGFβ1 mediates pro-fibrotic effects in EPCs of the co-culture system through a novel transcriptional complex including ph-Ezh2/Actin/Pol2. **(a)** Representative western blot analysis and quantification show a reduction in ph-EZH2 levels in EZH2-depleted EPCs versus control cells upon TGFβ1 stimulation for 72 h (n = 4 independent experiments, mean + s.d., *p < 0.05, Kruskal-Wallis/Dunn’s). **(b)** qPCR analysis of pro-fibrotic/EMT target genes in EZH2-depleted EPCs in co-culture with NHLFs shows that EZH2 is required for the effect of TGFβ1 on the upregulation of pro-fibrotic/EMT target genes (n = 4 independent experiments with 4 NHLFs donors, mean + s.d., ****p < 0.001, *p < 0.05, n = 4 independent experiments, ANOVA/Tukey’s). **(c)** ELISA analysis of signature FMT markers from NHLFs in the co-culture system shows that EHZ2-depleted EPCs prevent the effect of TGFβ1on the epithelial secretions driven FMT process (n = 4 NHLFs donors, mean + s.d., ****p < 0.001, **p = 0.004, *p = 0.0459, ANOVA/Tukey’s). **(d)** Representative western blot analysis of EZH2 co-immunoprecipitates shows increased levels of EZH2-bound POL2, ph-EHZ2, and actin in TGFβ1-treated EPCs. Unspecific IgG binding was used as a negative control. A representative from 3 independent experiments is shown. **(e)** Representative western blot analysis of POL2 immunoprecipitates shows co-precipitation of ph-EZH2 and nuclear actin. Increased levels of POL2-bound ph-EZH2 and nuclear actin are observed upon in EPCs subjected to apical TGFβ1 for 72 h. Unspecific IgG binding was used as a negative control. **(f)** ChIP-qPCR shows increased occupancy of POL2(S5p) at promoters of pro-fibrotic genes in EPCs subjected to TGFβ1 for 24 h. Negative IgG control is shown in Supplementary Fig. 3 (n = 3 independent experiments). **(g)** ChIP-qPCR shows increased occupancy of POL2(S2p) at pro-fibrotic genes in EPCs subjected to TGFβ1 for 48 h. Negative IgG control is shown in Supplementary Fig. 3 (n = 3 independent experiments).

To gain insights into the gene regulatory pathways responsible for this biological consequence, we performed gene set enrichment analysis (GSEA)^18^ and found that genes marked with H3K4me3 or H3K27me3^19^, as well as known target genes of the Polycomb subunits BMI1^20^, EZH2^21^ and SUZ12^22^ were significantly represented across all pairwise comparisons (Fig. 1n,o, Supplementary Fig. 1h and Supplementary Table 4). Importantly, GSEA also revealed a significant correlation with the IPF transcriptional signature^23,24^ in the TGFβ1-treated co-culture system (Fig. 1p and Supplementary Fig. 1k,l), confirming the relevance of our *in vitro* co-culture model to human disease.

Taken together, these observations suggest that TGFβ1-injured EPCs in the co-culture mediate their fibrotic effects through Polycomb activity. To test this hypothesis we measured global levels of H3K27me3 and EZH2 using immunofluorescence staining. A reduction in H3K27me3 was found in TGFβ1-damaged epithelium, whereas total EZH2 levels increased (Fig. 2a). We next determined H3K27me3 and EZH2 occupancy on pro-fibrotic genes detected in our RNA-seq and secretomics data, using chromatin immunoprecipitation (ChIP). We observed a decrease in H3K27me3 occupancy at promoters of pro-fibrotic genes in response to TGFβ1, consistent with their activation; however, EZH2 occupancy was increased (Fig. 2b,c).

The desynchronized occupancy of H3K27me3 and EZH2 prompted us to examine non-canonical functions of EZH2. Phosphorylation of EZH2 on T311^14^ or T487^13^ (ph-EZH2) has been shown to disrupt the interaction between EZH2 and SUZ12, a subunit required for PRC2 stabilization^25^. TGFβ1 treatment increased nuclear ph-EZH2 levels on both threonine sites (Fig. 2d and Supplementary Fig. 2), suggesting a PRC2-independent, non-canonical role of EZH2. We hypothesised that TGFβ1 leads to disruption of the EZH2-SUZ12 interaction, followed by a small reduction in global H3K27me3 levels. Since EZH1 has the ability to establish and maintain PRC2 function^26^, we sought to ascertain whether TGFβ1 stimulation leads to an EZ switch from EZH2-PRC2 to EZH1-PRC2, to sustain PRC2 activity. Co-Immunoprecipitation (co-IP) experiments uncovered an interaction between EZH1 and SUZ12 in which TGFβ1 enhanced EZH1-bound SUZ12, while simultaneously reducing EZH2-bound SUZ12 (Fig. 2e). To explore this EZ switch further at the single-gene level, we investigated EZH1, ph-EZH2 and H3K27me3 occupancy on pro-fibrotic genes and other known EZH2-PRC2 targets. ChIP analysis confirmed an enrichment of ph-EZH2 occupancy at the promoter regions of pro-fibrotic genes in response to TGFβ1 (Fig. 2f). As expected, EZH1 occupancy was increased at other non-fibrotic genes to maintain H3K27me3 levels (Fig. 2g,h), supporting previous reports of EZH1 compensation for EZH2 deficiency^27-29^.

As TGFβ1 leads to increased binding of ph-EZH2 at the promoters of pro-fibrotic genes, we analysed the contribution of ph-EZH2 in fine-tuning these genes. As expected, we observed an attenuation of ph-EZH2 upon EZH2 silencing in TGFβ1-treated EPCs (Fig. 3a). Subsequently, EZH2-depleted EPCs led to less pronounced upregulation of pro-fibrotic genes in response to TGFβ1 (Fig. 3b). This in turn prevented NHLFs from entering FMT in the co-culture system (Fig. 3c).

To unravel the mechanisms of the ph-EZH2-mediated pro-fibrotic gene expression, we investigated the potential binding partners of EZH2 upon TGFβ1-induced damage in EPCs. It has been shown that remodelling of the actin cytoskeleton is a major pathway involved in TGFβ1 signalling^30^. Actin is constantly shuttled between the cytoplasm and the nucleus, where it has been shown to associate with the chromatin remodelling complex and POL2 to enhance transcriptional activity^31-33^. As transcriptional regulation were among the most enriched GO term in TGFβ1-damaged EPCs, we hypothesized that ph-EZH2 may facilitate the transcriptional process of the pro-fibrotic genes through POL2 and nuclear actin. Intriguingly, analysis of the EZH2 co-IP upon TGFβ1 stimulation revealed an interaction between ph-EZH2, actin and POL2 (Fig. 3d). We further validated this novel interaction by performing reversed co-IP for the elongated form of POL2 phosphorylated on serine 2 (S2p)^34^ and observed that TGFβ1 promoted the formation of a novel fibrotic transcriptional complex consisting of ph-EZH2, POL2 and actin, as a consequence of the EZ switch. (Fig. 3e).

The emergence of an interaction between ph-EZH2 and POL2 led us to focus on the existence of poised and elongated forms of POL2^34^ at pro-fibrotic genes. Analysis of the occupancy of poised POL2 phosphorylated on Serine 5 (S5p) and elongated S2p revealed an increase in POL2(S5p) occupancy within the first 24 h upon TGFβ1 treatment, whereas POL2(S2p) occupancy only became visible after 48 h (Fig. 3f,g and Supplementary Fig. 3a-c). These data indicate that TGFβ1 induces a poised state of pro-fibrotic gene transcription, marked by POL2(S5p) and EZH2 interaction within the first 24 h. Such a poised state ensures only chronic and sustained TGFβ1 signals are capable of activating transcription through liberation of EZH2 from PRC2.

To understand how actin could impact the fibrotic transcriptional complex, we treated TGFβ1-induced pro-fibrotic EPCs with the Rho-kinase inhibitor Y27632^35^. Consistent with previous data^30^, TGFβ1 treatment increased actin polymerisation and non-muscle myosin II activity marked by the phosphorylation of its light chain (ph-MLC2, Fig. 4a,b). Furthermore, treatment with Y27632 but not EZH2 depletion can inhibit the TGFβ1-induced actomyosin remodelling. Importantly, disruption of actomyosin remodelling led to the abolition of the TGFβ1-induced fibrotic transcriptional complex, including ph-EZH2, POL2 and actin (Fig. 4c), suggesting that actin dynamics act as an upstream regulator of non-canonical EZH2.

**Figure 4.**
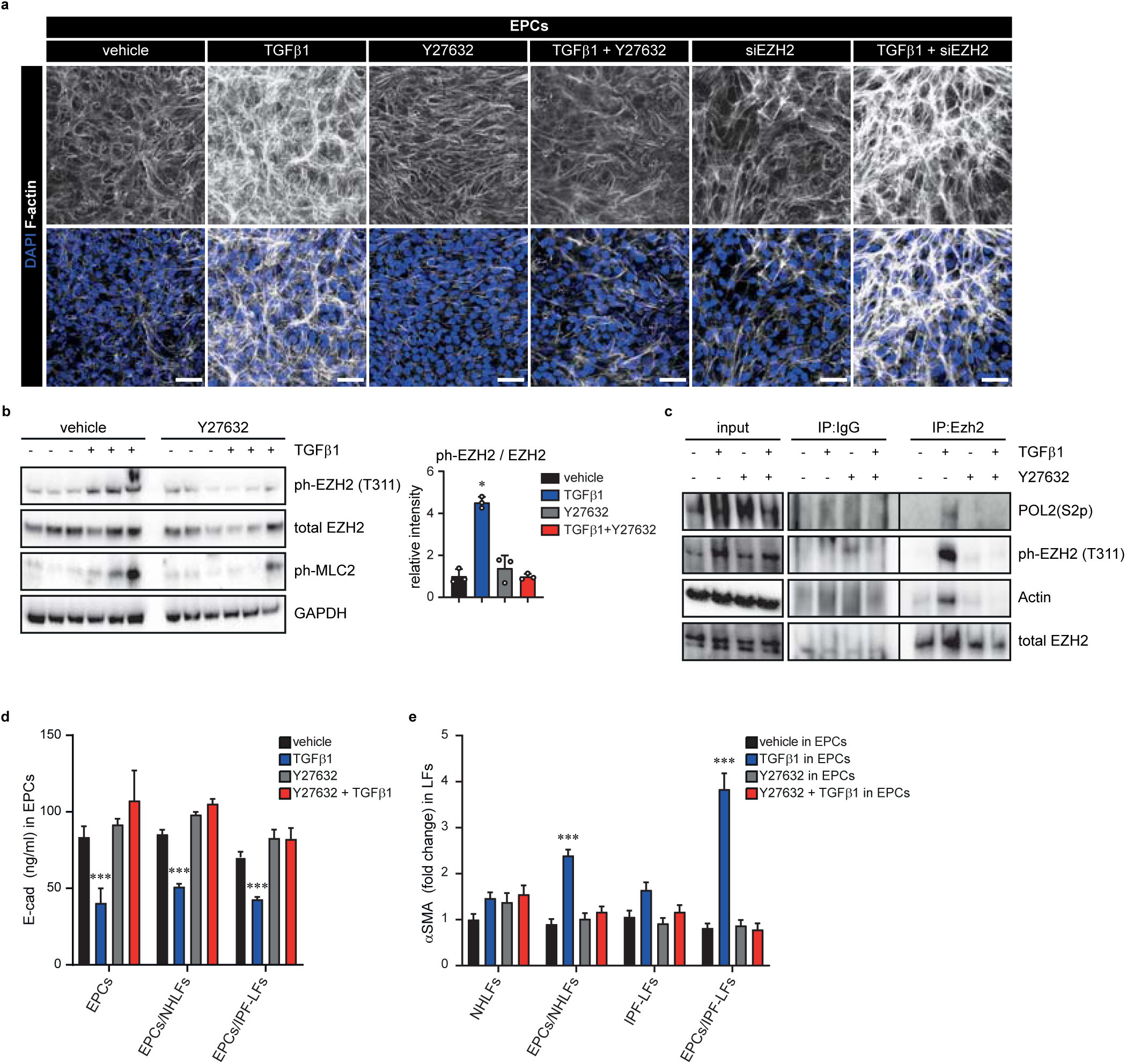
TGFβ1-mediated adjustment of fibrotic genes through actomyosin remodelling and EZH2. **(a)** Representative F-actin (phalloidin) and DAPI images of EPCs show that treatment with ROCK inhibitor Y27632 but not depletion of EZH2 can prevent TGFβ1-induced actomyosin remodelling in EPCs (scale bars 200μm). **(b)** Representative western blot analysis and quantification show that blocking actomyosin remodelling by Y27632 prevents TGFβ1-induced ph-EHZ2 in EPCs (n = 3 independent experiments, *p = 0.0523, Kruskal-Wallis/Dunn’s). **(c)** Representative immunoprecipates of EHZ2 shows abolition of TGFβ1-induced pro-fibrotic transcriptional complex of phEZH2-POL2-actin upon the convergent treatment of TGFβ1 and Y27632. Unspecific IgG binding was used as a negative control. Representative from 3 independent experiments is shown. **(d)** ELISA analysis of E-cad levels from EPCs in mono- or co-culture with NHLFs and IPF-LFs subjected to TGFβ1 for 72 h shows that blocking actomyosin remodelling by Y27632 prevents TGFβ1-reduced E-cad (n = 3 independent experiments with 6 NHLFs and 6 IPF-LFs donors, mean + s.d., ***p < 0.001, ANOVA/Tukey’s). **(e)** ELISA analysis of αSMA levels from NHLFs and IPF-LFs in mono and co-culture with EPCs shows that blocking actomyosin remodelling in EPCs leads to the abolition of epithelial secretions that drive increased αSMA upon TGFβ1 stimulation (n = 3 independent experiments with 6 NHLFs and 6 IPF-LFs donors, mean + s.d., ***p < 0.001, ANOVA/Tukey’s).

We next explored if the fibrotic crosstalk depended on epithelial actomyosin remodelling. To this end, we measured TGFβ1-induced pro-fibrotic proteins following inhibition of actomyosin remodelling by Y27632 and observed abolition of pro-fibrotic markers in both compartments of the co-culture system (Fig. 4d,e and Supplementary Fig 4). These data indicate that TGFβ1-injured EPCs trigger a fibrotic cascade with fibroblasts mediated by actomyosin remodelling.

To further validate our findings in a more physiological condition of human lung epithelium, we employed an air-liquid interface culture of primary stratified human lung small airway epithelial cells (SAECs) for 28 days and exposed this system to chronic TGFβ1 treatment during the last week of differentiation (Supplementary Fig. 5a). Upon chronic TGFβ1 treatment (tSAECs), we observed an abnormally heterogeneous epithelial layer compared to healthy SAECs (nSAECs) (Fig. 5a and Supplementary Fig. 5b), with no major defects in ciliation (Supplementary Fig. 5c). Importantly, nanoindentation measurements on tSAECs showed increased stiffness of the epithelium (Fig. 5b), within the range of a fibrotic lung^36^. This was accompanied by an increase in pro-fibrotic secretory proteins and loss of epithelial integrity (Fig. 5c and Supplementary Fig. 5d), demonstrating that our tSAECs model recapitulates major hallmarks of the IPF epithelium^4^.

**Figure 5.**
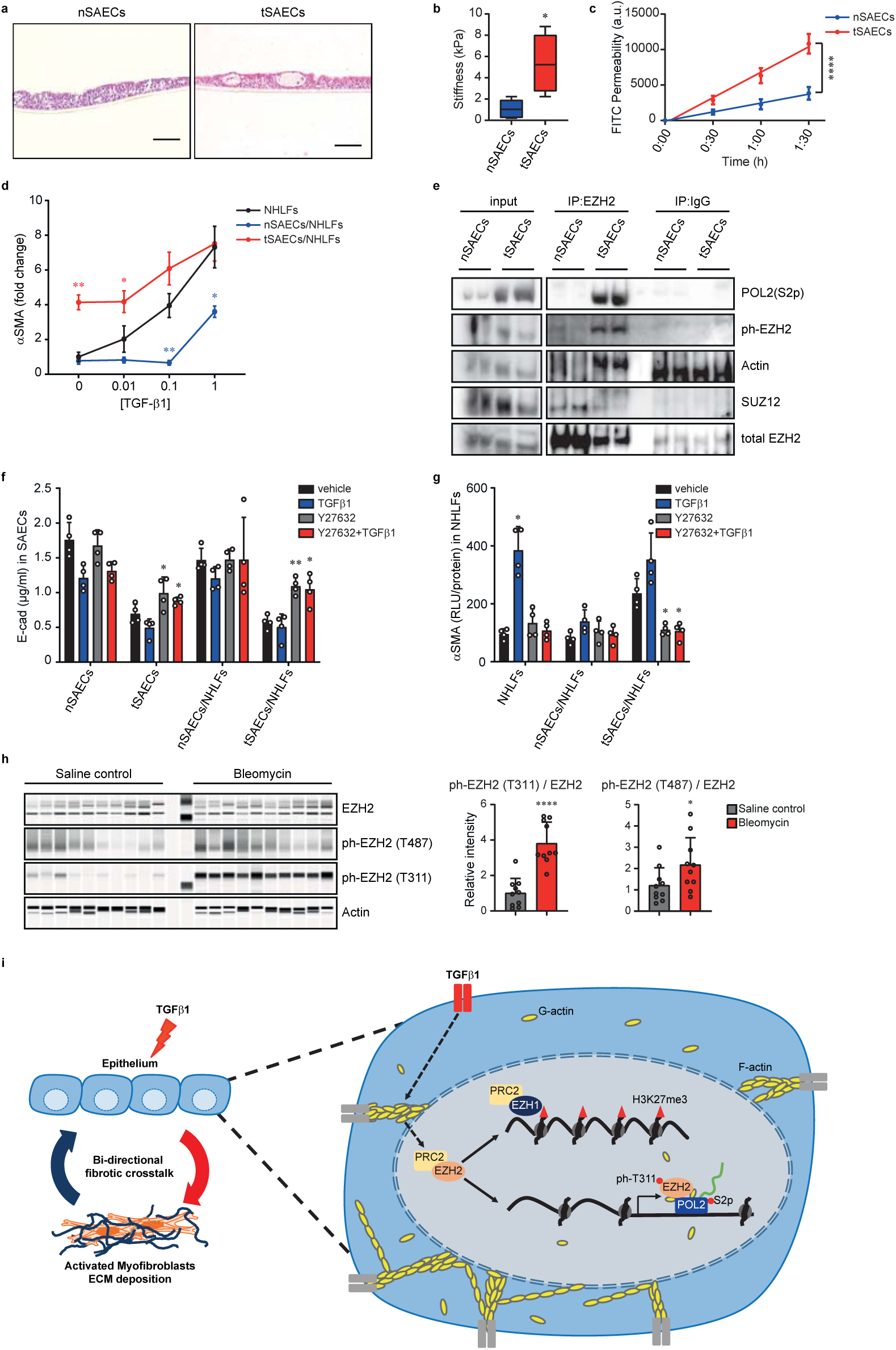
Non-canonical EZH2 as a potential therapeutic target in pulmonary fibrosis and its proposed mechanism. **(a)** Haematoxylin-eosin (H&E) staining of stratified and differentiated SAECs (scale bars 50μm). **(b)** Nanoindentation shows increased stiffness in tSAECs. Box plots display minimum, first quartile, median, third quartile and maximum (n = 5 SAECs donors, *p = 0.0179, two-sided paired t-test). **(c)** FITC dextran permeability assay reveals loss of epithelial integrity in tSAECs (n = 4 SAECs donors, mean ± s.d., ****p < 0.0001, Linear Regression). **(d)** ELISA analysis of αSMA levels from NHLFs in mono-(black line) or co-culture with nSAECs (blue) or tSAECs (red) shows an anti-fibrotic effect on nSAECs subjected to apical TGFβ1 for 72 h (n = 5 SAECs and 5 NHLFs donors,*p < 0.05, **p = 0.01, 2-way ANOVA /Tukey’s). **(e)** Representative western blot analysis of EZH2 immunoprecipitates shows co-precipitation of ph-EZH2, POL2(S2p) and actin in tSAECs. Note the loss of EZH2-bound SUZ12 in tSAECs compared to nSAECs. Unspecific IgG was used as a negative control. **(f)** ELISA analysis shows decreased E-cad levels in tSAECs in the mono- or co-culture system. This reduction can be rescued by the ROCK inhibitor Y27632 (n = 4 SAECs donors, mean + s.d., *p < 0.05, **p = 0.0052, 2-way ANOVA/Tukey’s). **(g)** ELISA analysis of aSMA levels shows increased aSMA levels in NHLFs when co-cultured with tSAECs. Blocking actomyosin remodelling by Y27632 can prevent this effect (n = 4 NHLFs donors, mean + s.d., *p < 0.05, 2-way ANOVA/Tukey’s). **(h)** Representative western blot analysis and quantification show increased ph-EZH2 levels in Bleomycin treated mice compared to saline control group (n = 10 mice / treatment, mean + s.d., *p = 0.04, ****p < 0.001, Mann-Whitney). **(i)** A model describing TGFβ1-injured epithelium initiating a bi-directional fibrotic crosstalk with fibroblasts. TGFβ1-injured epithelium promotes (1) an EZ-switch from EZH2-PRC2 to EZH1-PRC1, which is required to maintain H3K27me3 at TGFβ1 non-target genes; and (2) a PRC2-independent EZH2 that forms a pro-fibrotic transcriptional complex with POL2(S2p) and nuclear actin to fine-tune transcription at pro-fibrotic genes.

We next examined the pro-fibrotic potential of this stiffened epithelium in co-culture with LFs. As expected, tSAECs activated the fibrotic cascade in the co-culture system without further TGFβ1 treatment, whereas nSAECs exhibited anti-fibrotic effects in the presence of low dose TGFβ1 (Fig. 5d and Supplementary Fig. 5e), confirming a vital role of epithelial cells in the pathogenesis of IPF.

In addition to the pro-fibrotic phenotype of tSAECs, we observed a dramatic elevation of ph-EZH2 levels and the formation of the previously described ph-EZH2 transcription complex (Fig. 5e). As expected, the EZH2-SUZ12 interaction as part of the PRC2 complex was lost in tSAECs. Furthermore, acute Y27632 treatment reduced the pro-fibrotic effects in tSAECs (Fig. 5f,g), validating the role of actomyosin remodelling in the fibrotic cascade.

Finally, to assess the non-canonical EZH2 pathology *in vivo*, we analysed a Bleomycin-induced pulmonary fibrosis mouse model. Lung lysates were examined at day 21 after bleomycin or saline instillation. Consistent with our *in vitro* data, we detected increased ph-EZH2 levels in bleomycin-induced pulmonary fibrosis compared to those of saline controls (Fig. 5h). This result demonstrates that the molecular mechanisms described here are ubiquitous in different fibrotic model systems and are recapitulated in whole organ fibrosis.

Our study provides direct evidence that epithelial injury is a potent stimulus for the initiation of a fibrotic cascade, as summarized in Fig. 5i. While earlier reports have suggested a vital role of EZH2 in fibrotic progression^10,37^, little is known about its precise mechanisms. We propose a novel fibrotic model in which TGFβ1-injured lung epithelium promotes two mechanisms: (1) an EZ-switch from EZH2-PRC2 to EZH1-PRC2, which is required to maintain silencing at non-fibrotic genes; and (2) a PRC2-independent role of EZH2 that forms a pro-fibrotic transcriptional complex including ph-EZH2, POL2 and nuclear actin to fine-tune transcription.

POL2(S5p) has been reported to bind to EZH2-PRC2 at lineage-specific genes. This interaction establishes poised promoters that alter the chromatin state to ensure only strong and sustained signals are capable of driving transcription^38,39^. In our model, this process of transcriptional activation occurs in a step-wise fashion. Within the first 24 h of TGFβ1 treatment, the establishment of poised promoters at pro-fibrotic genes is induced. Consequently, accessibility to these genes is gradually increased, driving EPCs to a plastic and inducible epigenetic state. Chronic exposure to TGFβ1 will then lead to loss of PRC2 occupancy at these sites. Loss of repressive complexes has been suggested to lower chromatin thresholds for aberrant gene activation^38^, similar to the here reported liberation of EZH2 coupled with an EZ-switch. Importantly, we show an interaction between PRC2-independent EZH2 and the elongated form of POL2 to mediate pro-fibrotic gene expression. This is an intriguing new function of EZH2 as a transcriptional activator. Further studies should be directed towards better understanding of this novel interaction between EZH2 and POL2(S2p).

Taken together, our study sheds new light on the non-canonical functions of EZH2 as a transcriptional activator and thereby implies that enzymatic inhibitors of EZH2 may not be efficacious unless they are capable of disrupting its interaction with the transcriptional machinery, while maintaining H3K27me3 occupancy at non-target genes. Since current epigenetic inhibitors lack the ability to modify epigenetic states in a precise manner and the efficiency to target specific genomic loci^40^, we have identified a novel pro-fibrotic mechanism that could serve as a potential therapeutic target for the next generation of epigenetics therapeutics in IPF treatment.

## ACKNOWLEDGEMENTS

We thank S. A. Wickström, J. Morgner and members of lung repair and regeneration department for critical reading of the manuscript, M. Kästle for sharing the whole lung lysate from bleomycin-induced fibrosis mouse model, J. Li, D. Veyel and A. R. Campos for supporting on proteomics, J. Schütt, C. Keller and E. Peter for technical assistance, and the Global Computational Biology Department, Boehringer Ingelheim Pharma GmbH & Co. KG for sequencing. This work is funded by Boehringer Ingelheim Pharma GmbH & Co. KG and UK Medical Research Foundation Fellowship (MRF-091-0001-RG-GARNE to J.P.G.).

## AUTHOR CONTRIBUTIONS

H.Q.L. and J.P.G. conceived and supervised the study. H.Q.L. and M.A.H. designed, performed and analysed most of the experiments. H.Q.L. performed the bioinformatics analyses. K.Q. assisted with the RNA-seq. I.K., W.S., V.S., J.W., and E.S. performed experiments. J.P.G., F.E.H. and M.J.T. assisted with initial design and concept for the co-culture model. H.Q.L. and J.P.G. provided conceptual advice. H.Q.L. and M.A.H. wrote the paper. All authors commented on and edited the manuscript.

## Materials and methods

### Primary cells and cell lines

Human lung fibroblasts from healthy (NHLFs, Lonza, CC-2512) and IPF (IPF-LFs, Lonza, CC-7231) were grown in Fibroblast Growth Medium-2 (GIBCO, CC-3132). Fibroblasts from passages 4-8 were used for all experiments. A549 alveolar epithelial cells (EPCs, Sigma-Aldrich, 86012804) were grown in DMEM with 10% fetal calf serum (GIBCO). Human small airway epithelial cells (SAECs, Lonza, CC-2547) were cultured in PneumaCult-Ex Plus medium (Stemcell Technologies, 05040). SAECs were used between passages 2-4.

### EPCs/LFs co-culture

200 000 EPCs were seeded on 24 mm Transwell (Corning, 0.4 µm pore size, PET membrane, 3450). 100 000 Fibroblasts per well (6-well plate, Corning, 3516) were seeded. Cells were starved overnight and medium was replaced by fresh DMEM with 200 µl apical and 2000 µl basolateral before experiment start. The two compartments were then brought together and treatments were applied apically for 72 h.

### SAECs/LFs co-culture

300 000 SAECs were seeded on collagen type I (Corning) coated 24 mm Transwell (Corning, 0.4 µm pore size, PET membrane, 3450) in PneumaCult-Ex Plus medium until confluent. Apical medium was then removed and SAECs were stratified and differentiated at air-liquid interface in Pneumacult-ALI-S medium (Stemcell Technologies, 05099) for 28 days. The basolateral medium was exchanged every other day. Where indicated, chronic TGFβ1 treatment was applied in basolateral medium after 14 days post ALI (dpa), when cilia were detected. After full stratification and differentiation (28 dpa), SAECs were brought together with fibroblasts and treatments were applied basolaterally.

### RNA sequencing analysis and bioinformatics

After 72 h of TGFβ1 treatment in the co-culture, transcriptional profiles of both EPCs and NHLFs/IPF LFs were analysed with RNA sequencing. Total RNA was extracted and purified using RNeasy Plus Mini Kit (QIAGEN) according to the manufacturer’s instructions. Quantification and quality control of total RNA was measured with an RNA Pico chip on a Bioanalyzer (Agilent, Santa Clara, CA, US), followed by adjustment of total RNA to equal amounts. From this RNA, libraries for sequencing were prepared from using the TrueSeq RNA Sample Prep Kit v2-Set B (Illumina, San Diego, CA, US) according to the manufacturer’s instructions. Single-end sequencing was performed on an Illumina HiSeq2000 instrument using the TruSeq SBS Kit HS-v3 (50-cycle) (Illumina, San Diego, CA, US).

The processing pipeline was carried out as previously described ^1^. Differentially expressed gene analysis was done using the limma R-package with Benjamini-Hochberg correction^2^. Genes with an adjusted *P* value < 0.05 were considered significant.

Gene ontology term analyses were performed using Enrichr^3^. Gene Set Enrichment Analysis was performed on a pre-ranked gene list according to log_2_ fold change and compared with the Board Institute Molecular Database collection of chemical and genetic perturbations (C2 CGP, 3297 gene sets) or Oncogenic signatures (C6, 189 gene sets) using the web-based tool available from the Broad Institute^4,5^.

### Mass-Spectrometry

#### Sample preparation

Supernatant medium was buffer exchanged using 3.5-kDa Amicon filter (Millipore) to 8 M urea, 50 mM ammonium bicarbonate buffer. While in the filter, proteins were reduced with 5 mM tris(2-carboxyethyl)phosphine (TCEP) at 30°C for 60 min, and subsequently alkylated with 15 mM iodoacetamide (IAA) in the dark at room temperature for 30 min. The buffer was then exchanged again to 1 M urea, 50 mM ammonium bicarbonate, the sample was recovered from the Amicon tube into a new microfuge tube and protein concentration was determined using bicinchoninic acid (BCA) protein assay (Thermo Scientific). Proteins were subjected to overnight digestion with mass spec grade Trypsin/Lys-C mix (1:25 enzyme/substrate ratio). Following digestion, samples were acidified with formic acid (FA) and subsequently desalted using AssayMap C18 cartridges mounted on an Agilent AssayMap BRAVO liquid handling system. Cartridges were sequentially conditioned with 100% acetonitrile (ACN) and 0.1% FA, samples were then loaded, washed with 0.1% FA, and peptides eluted with 60% ACN, 0.1% FA. Finally, the organic solvent was removed in a SpeedVac concentrator prior to LC-MS/MS analysis.

#### Mass spectrometry

Prior to LC-MS/MS analysis, dried peptides were reconstituted with 2% ACN, 0.1% FA and concentration was determined using a NanoDropTM spectrophometer (ThermoFisher). Samples were then analyzed by LC-MS/MS using a Proxeon EASY-nanoLC system (ThermoFisher) coupled to an Orbitrap Fusion Lumos mass spectrometer (Thermo Fisher Scientific). Peptides were separated using an analytical C18 Acclaim PepMap column (75µm x 500 mm, 2µm particles; Thermo Scientific) at a flow rate of 300 nL/min (60° C) using a 75-min gradient: 1% to 5% B in 1 min, 6% to 23% B in 44 min, 23% to 34% B in 28 min, and 34% to 48% B in 2 min (A= FA 0.1%; B=80% ACN: 0.1% FA). The mass spectrometer was operated in positive data-dependent acquisition mode. MS1 spectra were measured in the Orbitrap in a mass-to-charge (m/z) of 375 – 1500 with a resolution of 60,000 at m/z 200. Automatic gain control target was set to 4 x 105 with a maximum injection time of 50 ms. The instrument was set to run in top speed mode with 2-second cycles for the survey and the MS/MS scans. After a survey scan, the most abundant precursors (with charge state between +2 and +7) were isolated in the quadrupole with an isolation window of 1.6 m/z and fragmented with HCD at 30% normalized collision energy. Fragmented precursors were detected in the ion trap as rapid scan mode with automatic gain control target set to 1 x 104 and a maximum injection time set at 35 ms. The dynamic exclusion was set to 20 seconds with a 10 ppm mass tolerance around the precursor.

#### MS data processing

All raw files were processed with MaxQuant^6^ (version 1.5.5.1) using the integrated Andromeda Search engine^7^ against a target/decoy version of the curated human Uniprot proteome without isoforms (downloaded in January of 2019) and the GPM cRAP sequences (commonly known protein contaminants). First search peptide tolerance was set to 20 ppm, main search peptide tolerance was set to 4.5 ppm. Fragment mass tolerance was set to 20 ppm. Trypsin was set as enzyme in specific mode and up to two missed cleavages was allowed. Carbamidomethylation of cysteine was specified as fixed modification and protein N-terminal acetylation and oxidation of methionine were considered variable modifications. The target-decoy-based false discovery rate (FDR) filter for spectrum and protein identification was set to 1%.

#### Data analysis

Quantitative analysis of the proteome data was performed in the R statistical programming language (version 3.5.1, 2018-07-02) using in-house R script wrapper for Bioconductor packages such as limma and MSstats. Briefly, feature (a peptide sequence of a given charge state and with potential amino acid modifications) intensities were log2-transformed and loess-normalized to account for systematic errors. Testing for differential abundance was performed using MSstats bioconductor package based on a linear mixed-effects model. In cases where a protein was completely missing in one of the conditions being compared, we imputed an empirical log_2_ fold change and *P* value. Imputed log_2_ fold change was calculated by summing up the intensities of the protein across the replicates in the condition where it was detected, then dividing it by 3.3 and taking the logarithm base 2 of the number, whereas its imputed pvalue was calculated by dividing 0.05 by the number of replicates the protein was detected.

### Chemical treatments

Where indicated, cells were treated with ROCK inhibitor (Y27632, Sigma-Aldrich, 10 µM), EZH2 inhibitor (GSK126, 10μnM) and TGFβ1 (R&D, 240-B, 1 ng/ml). The vehicle dimethylasulfoxide (DMSO) was used as control for Y27632 and GSK126 treatments, 4mM HCl (Sigma-Aldrich) containing 1 mg/ml BSA (Sigma-Aldrich) was used as control for TGFβ1 treatment.

### RNAi transfection

siRNA targeting hEZH2 (ID: s4916 and s4918) and negative control siRNA (AM4635) were from Thermo Fisher Scientific (Silencer Select). Transfection were performed 1 day before co-culture in EPCs at 70% confluence, using Lipofectamine RNAiMAX (Invitrogen) according to manufecturer’s instruction. After 24 h of transfection, EPCs were subjected to co-culture experiment.

### Co-immunoprecipitation (co-IP) and western blot

#### Co-Immunoprecipitation

Co-IP was performed using following antibodies: EZH2 (Active Motif, 39933, 1:100), Pol2-S2p (Active Motif, 61083, 1:100), SUZ12 (Cell Signalling, 3737, 1:100). Cells were harvested in IP lysis buffer (Thermo Fisher Scientific, 87788), containing protease and phosphatase inhibitors. Lysates were cleared by centrifugation at 10, 000 x g and protein concentration was quantified with Pierce BCA (Thermo Fisher Scientific, 23225).

Antibodies were coupled to Protein A/G magnetic beads (Thermo Fisher Scientific, 26162) for 1 h at 4°C. Lysates were then incubated with 50 μl beads-antibodies complex overnight at 4°C. An isotype IgG antibody (Cell Signaling, 2729S) was used as the control. After repeated washing, proteins were eluted in NuPAGE LDS Sample buffer (Novex, NP0007) at 70°C in 10 min, followed by western blot analysis.

#### Western Blotting

Western blot was carried out as previously described^8^. The following antibodies were used: β-Actin (Cell Signalling, 3700, 1:5000), EZH1 (Cell Signalling, 42088, 1:5000), EZH2 (Cell Signalling, 5246, 1:5000), EZH2 (Cell Signalling, 3147, 1:5000), GAPDH (Cell Signalling, 3683, 1:100000), H3K27m3 (Cell Signalling, 9733, 1:5000), ph-EZH2 (T487) (Invitrogen, PA5-105660, 1:1000), ph-EZH2 (T311) (Cell Signalling, 27888, 1:1000), Phospho-Myosin Light Chain 2 (Thr18/Ser19) (Cell Signalling, 3674, 1:1000), POL2-S2p (Active Motif, 61083, 1:1000), POL2-S5p (Cell Signalling, 13523, 1:1000), POL2 (Cell Signalling, 2629, 1:1000), SUZ12 (Cell Signalling, 3737, 1:1000).

### RT-qPCR

RNA was isolated using the RNAeasy Plus Mini Kit (QIAGEN), after which reverse transcription was performed using the High-Capacity cDNA Reverse Transcription Kit (Bio-rad). qPCR was performed on the ABI ViiA 7 real-time PCR System (Life Technologies) using QuantiFast Probe PCR kit master mix (QIAGEN) with FAM- or VIC-labelled TaqMan MGB probe (Applied Biosystem). Changes in gene expression were calculated using the comparative cycle threshold method with HPRT1, GAPDH or S26 as house-keeping controls. For a complete list of all probes used in this study see Supplementary Table 5a.

### Chromatin immunoprecipitation (ChIP) analyses

ChIP analyses were carried out as described previously^8^. In brief, cells were crosslinked in 1% formaldehyde (Sigma-Aldrich), after which cells were lysed and sonicated to fragment DNA. Following sonication, 10% was taken for input and lysates were incubated with 5 µg antibody or isotype IgG overnight at 4°C. The next day, lysates were incubated with protein A/G beads (Dynabeads, Life Technologies) followed by extensive washing. Chromatin was then decrosslinked and eluted. DNA was purified and analysed by qPCR. ChIP-qPCR was performed using DyNamo Color Flash SYBR Green Mix (Thermo Fisher). Enrichment was determined by normalising to input DNA of the target gene as a percentage of input. For a complete list of primers see Supplementary Table 5b.

The following antibodies were used: EZH1 (Cell Signalling, 42088) EZH2 (Active Motif, 39933), H3K27m3 (Cell Signalling, 9733), POL2-S2p (Active Motif, 61083), POL2-S5p (Cell Signalling, 13523), ph-EZH2 (T487) (Invitrogen, PA5-105660), ph-EZH2 (Thr311) (Cell Signalling, 27888).

### ELISA

The supernatant from the basolateral compartment of mono- and co-cultures were harvested and MMP7 (Meso Scale Discovery (MSD), F210K), MCP-1 (MSD, K151UGK), Fibronectin (Thermo Fisher Scientific, BMS2028TEN), TGFβ1 (R&D, DY240), and COL1A1 (R&D, DY6220) were analysed according to the manufacturer’s instructions. Cell lysates were extracted in RIPA buffer (Sigma-Aldrich, R0278) and E-cadherin (MSD, F21YX) or α-SMA (custom made, high binding plate from MSD, L15XB, with α-SMA antibody from Sigma-Aldrich, A2547) were measured in accordance with the manufacturer’s instructions. ELISA signal was measured either on a MESO SECTOR® 6000 using the MSD Discovery Workbench Software 4.0 (MSD, LLC., Rockville, MD, USA) or on a SpectraMaxM5 spectrophotometer (Molecular Devices) using SoftMax Pro 6.5 Software. Total protein concentration of the supernatant measured using the Pierce BCA protein assay kit (Thermo Scientific) was used as a control.

### Immunofluorescence and confocal microscopy

Cells were were fixed in 4% paraformaldehyde or ice-cold methanol, permeabilised with 0.3% Triton X-100 in PBS, and blocked in 5% bovine serum albumin (BSA). Samples were incubated overnight in primary antibody followed by washing and incubation in secondary antibody and/or phalloidin labelling. Finally, samples were mounted with Prolong Antifade Mountant with DAPI (Invitrogen P36935) and coversliped. The following primary antibodies were used: α-SMA (Sigma-Aldrich A2547; 1:500), E-cadherin (BD 310181; 1:500), Collagen type I (Sigma-Aldrich SAB4200678; 1:300), H3K27me3 (Cell Signalling 9733; 1:500), EZH2 (Cell Signalling 3147; 1:500), ph-EZH2 T311 (Cell Signalling 27888; 1:500).

All fluorescence images were collected by laser scanning confocal microscopy (LSM710; Zeiss) with an AXIO Observer Z1 using 40x or 63x immersion objectives. All image analyses was carried out using ImageJ software.

### Histology

Haematoxylin-eosin staining of paraffin-embedded SAECs sections was performed using standard protocols. Images were collected with a Zeiss Axio Observer system.

### 10S FITC-Dextran permeability assay

Following 4 week differentiation, permeability of the epithelial layer was monitored by transcellular passage of 10 kDa dextran labelled with fluorescein isothiocyanate (FITC). Growing medium was substituted with RPMI without phenol red (Gibco). 5 mg/ml of 10 µl FITC-Dextran was applied to the apical side of the cells and was incubated at 37°C, 5% CO_2_. Passage of FITC dextran was measured at 30 min, 60 min and 90 min intervals, using a SpectraMaxM5 spectrophotometer (Molecular Devices) and SoftMax Pro 6.5 Software.

### Force indentation spectroscopy

Nanoindentation was performed on airlifted SAECs layer using a Pavone nanoindenter (Optics 11, Netherlands). Indenter probes had a glass spherical tip (diameter ∼30 μm) mounted on an individually calibrated cantilever with a spring constant of ∼0.025 N m^−1^. For all indentation, forces of up to 3 nN were applied and the velocities of cantilever approach and retraction were kept constant at 2 µm s^-1^ ensuring detection of elastic properties only. The effective Young’s modulus was calculated using a linear Hertzian contact model based on the first 10% of the force–distance curve. For each sample, five areas were measure of 50 x 50 µm.

### Statistical analysis and reproducibility

Statistical analyses were performed using GraphPad Prism (version 8.0). Statistical significance was determined by the Mann-Whitney U-test, Wilcoxon test, paired t-test, Kruskal-Wallis ANOVA with Dunn’s, Tukey’s or Dunnett’s post hoc test, non-linear / linear regression or Spearman’s rank correlation coefficient test as indicated in the corresponding figure legends. In all comparisons which a test for normally distributed data was used, Gaussian distribution was determined with Kolmogorov-Smirnov test (α = 0.05).

All experiments in this study were repeated for at least three independent times / biological replicates.

### Data availability

RNA-seq data that support this study have been deposited in the Gene Expression Omnibus (GEO) and the secretomics data in the proteomics repository MassIVE and will be made public upon acceptance of the manuscript. All other data that support the finding are available from the authors on request.

---

**Supplementary Figure 1.**
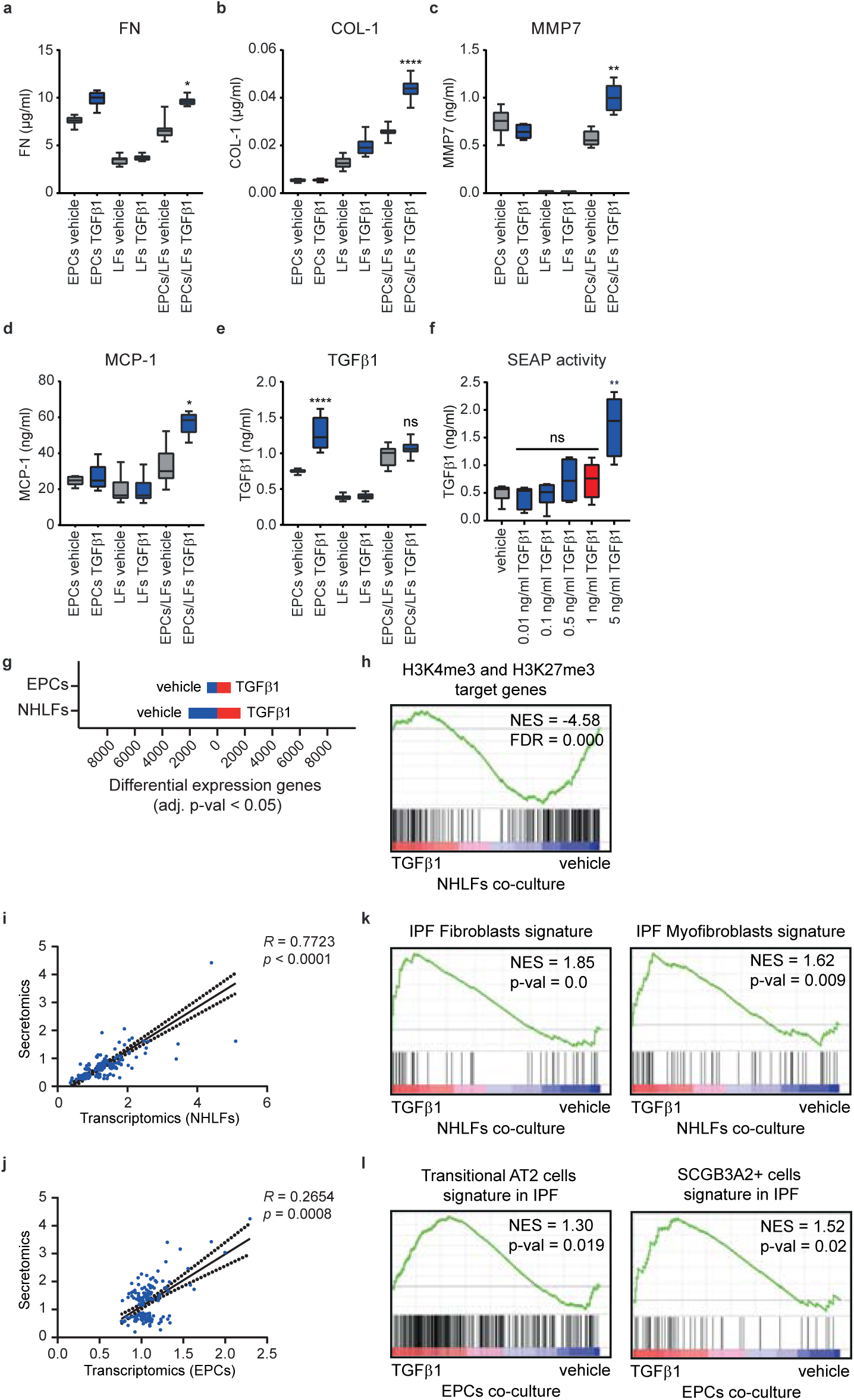
Characterisation of the EPCs/LFs co-culture system, related to Figure 1. **(a-e)** ELISA analysis of pro-fibrotic markers in the mono- and co-culture system. Note elevated levels of pro-fibrotic markers in the TGFβ1-treated co-culture after 72 h (n = 6 independent experiments, box plots (minimum, first quartile, median, third quartile and maximum), ****p < 0.001, *p < 0.05, **< 0.01, ANOVA/Tukey’s). **(f)** Secreted alkaline phosphatase protein (SEAP) assay shows detectable induction of TGFβ1 in basolateral compartment only after treatment with 5 ng/ml TGFβ1 apically (n = 4 independent experiments, box plots (minimum, first quartile, median, third quartile and maximum), **p = 0.0084, ns = non-significant, Kruskal-Wallis). **(g)** Summary of DEG analysis from RNA-seq of both compartments from the co-culture systems. **(h)** GSEA shows enrichment of genes marked by H3K4me3 and H3K27me3 in TGFβ1-treated NHLFs from the co-culture system. **(i, j)** Scatter plots show a positive correlation between secreted protein (p-val < 0.05) and RNA-seq from NHLFs (i) or EPCs (j) from the co-culture system (R is Spearman’s rank correlation coefficient). **(k, l)** GSEA of NHLFs (k) and EPCs (l) from the co-culture system shows an enrichment of the IPF transcriptional and cellular phenotype in TGFβ1-treated EPCs/LFs co-culture system.

**Supplementary Figure 2.**
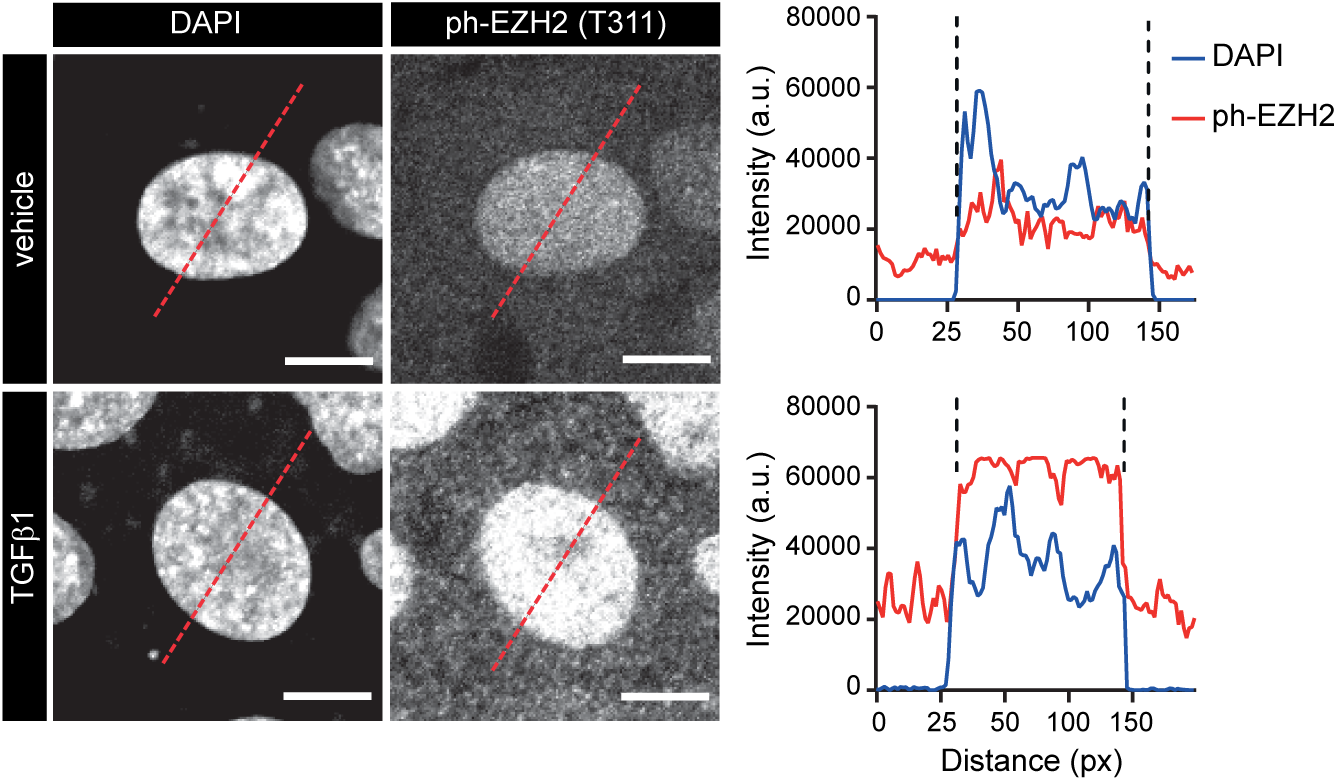
Induction of ph-EZH2 in EPCs upon TGFβ1 stimulation, related to Figure 2. Representative ph-EZH2 images of a confocal plane at the level of the nucleus from vehicle and TGFβ1-treated EPCs (scale bars 7.5μm). Right side panels show linescans through the cytoplasm and the nucleus indicating enrichment of ph-EZH2 in both compartments upon TGFβ1 stimulation. Representation from 3 independent experiments is shown.

**Supplementary Figure 3.**
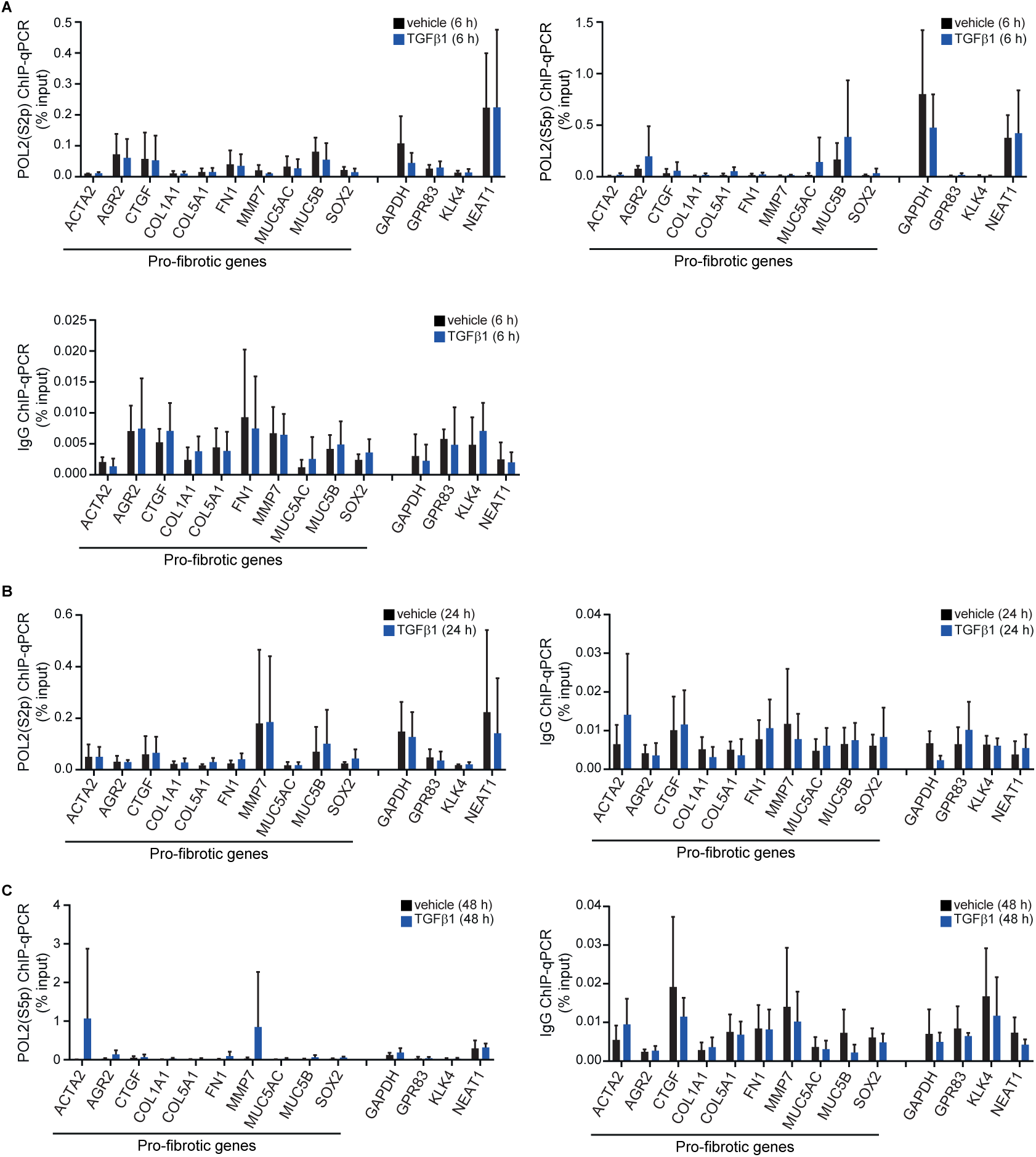
Analysis of transcriptional activation dynamics upon TGFβ1 treatment in EPCs, related to Figure 3. **(a)** ChIP-qPCR analysis of POL2(S2p) and POL2(S5p) in EPCs subjected to TGFβ1 for 6 h. Unspecific IgG was used as negative control (n = 3 independent experiments). **(b)** ChIP-qPCR analysis of POL2(S2p) in EPCs subjected to TGFβ1 for 24 h. Unspecific IgG was used as negative control (n = 3 independent experiments). **(c)** ChIP-qPCR analysis of POL2(S5p) in EPCs subjected to TGFβ1 for 48 h. Unspecific IgG was used as negative control (n = 3 independent experiments).

**Supplementary Figure 4.**
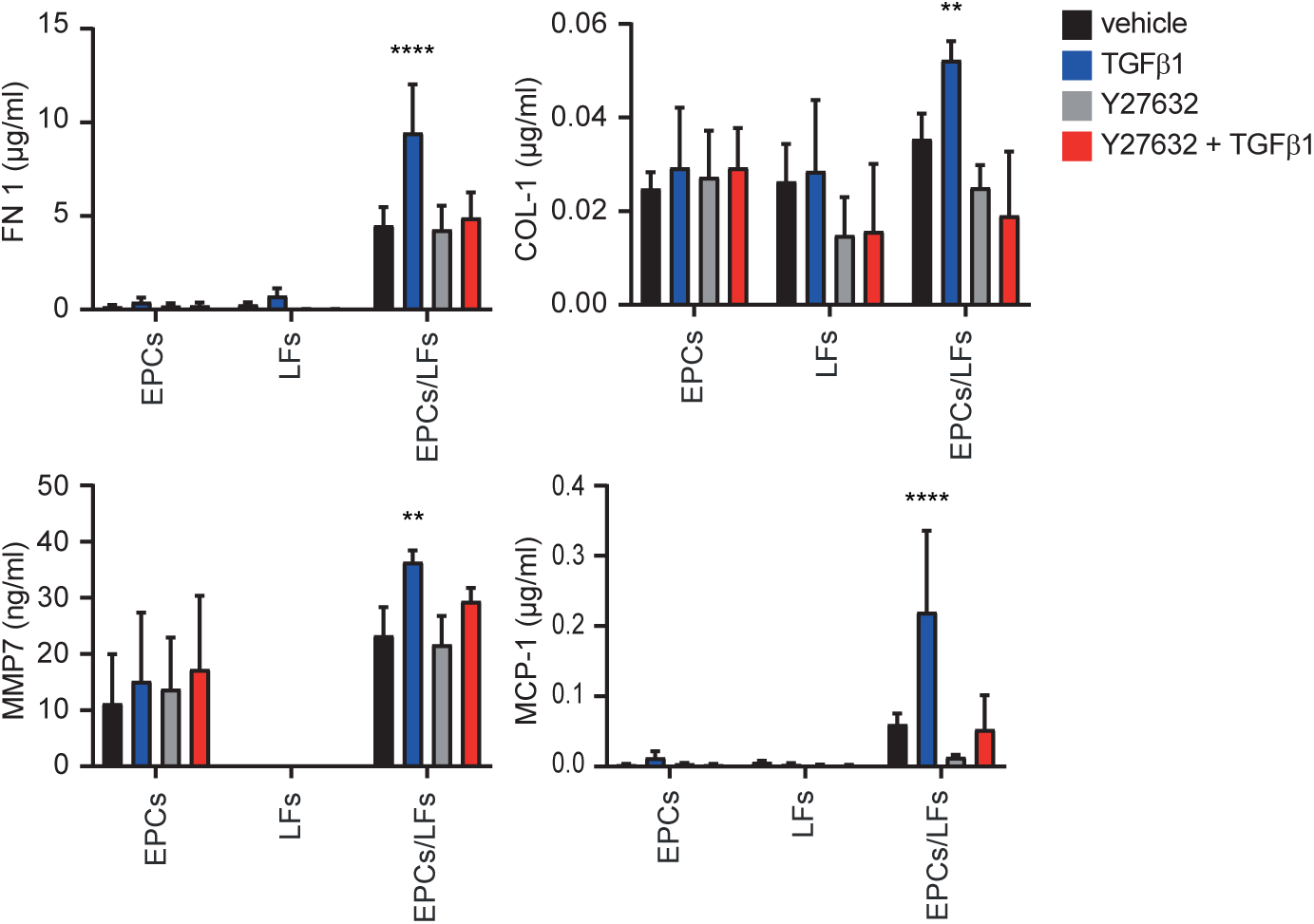
Induction of pro-fibrotic markers is driven by actomyosin remodeling, related to Figure 4. ELISA analysis of pro-fibrotic markers from the co-culture shows that blocking actomyosin remodelling by Y27632 prevents TGFβ1-induced elevation of FN1, COL-1, MMP7 and CP-1 (n = 3 independent experiments with 6 LFs donors, mean + s.d., **p < 0.01, ****p < 0.001, ANOVA/Tukey’s).

**Supplementary Figure 5.**
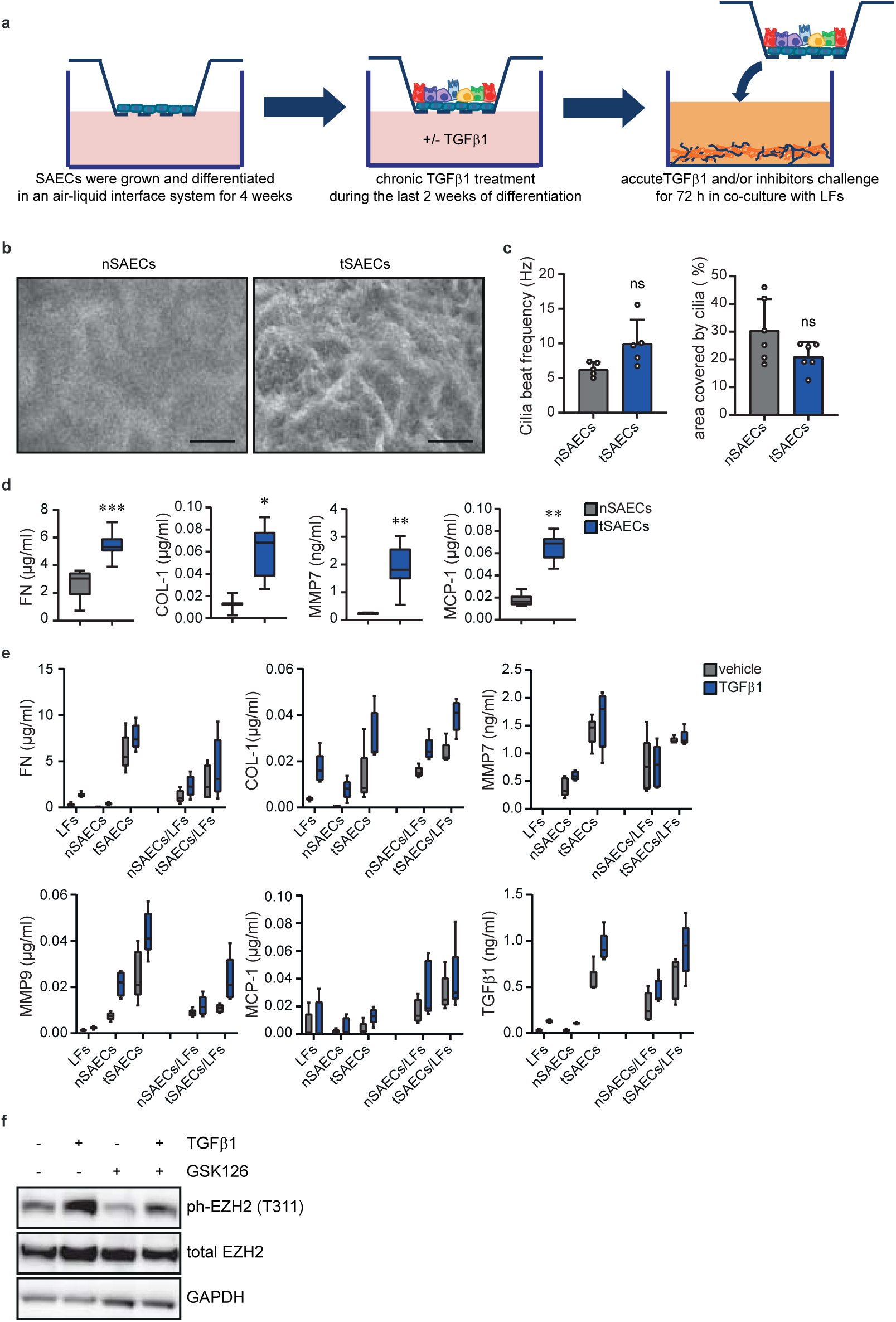
Characterisation of the SAECs/LFs co-culture system, related to Figure 5. **(a)** Schematic workflow of chronic TGFβ1 treatment during SAECs stratification and differentiation, followed by co-culture with LFs. **(b)** Representative images showing morphological changes of SAECs upon chronic TGFβ1 treatment (scale bars 500μm). **(c)** Analysis of cilia beat frequency and cilia area shows no significant differences between nSAECs and tSAECs (n = 3 donors, mean + s.d., ns = non-significant, paired t-test) **(d)** ELISA analysis shows increased pro-fibrotic proteins secretion in tSAECs (n = 3 independent experiments from 4 donors, *p = 0.0313, **p = 0.0078, ***p = 0.0002, paired Wilcoxon test, box plots show minimum, first quartile, median, third quartile and maximum). **(e)** ELISA analysis of pro-fibrotic markers in the mono- and co-culture system between SAECs and LFs. Note elevated pro-fibrotic levels in tSAECs and the co-culture system. (n = 3 independent experiments from 4 SAECs donors and 6 LFs donors, box plots show minimum, first quartile, median, third quartile and maximum). **(f)** Western blot analysis shows reduced ph-EZH2 levels upon GSK126 treatment in tSAECs.

## References

1 Zepp, J. A. & Morrisey, E. E. Cellular crosstalk in the development and regeneration of the respiratory system. Nat Rev Mol Cell Biol 20, 551–566, doi:10.1038/s41580-019-0141-3 (2019).

2 Sasai, Y. Cytosystems dynamics in self-organization of tissue architecture. Nature 493, 318–326, doi:10.1038/nature11859 (2013).

3 Wickstrom, S. A. & Niessen, C. M. Cell adhesion and mechanics as drivers of tissue organization and differentiation: local cues for large scale organization. Curr Opin Cell Biol 54, 89–97, doi:10.1016/j.ceb.2018.05.003 (2018).

4 Hewlett, J. C., Kropski, J. A. & Blackwell, T. S. Idiopathic pulmonary fibrosis: Epithelial-mesenchymal interactions and emerging therapeutic targets. Matrix Biol 71-72, 112–127, doi:10.1016/j.matbio.2018.03.021 (2018).

5 Sakai, N. & Tager, A. M. Fibrosis of two: Epithelial cell-fibroblast interactions in pulmonary fibrosis. Biochim Biophys Acta 1832, 911–921, doi:10.1016/j.bbadis.2013.03.001 (2013).

6 Selman, M. & Pardo, A. The leading role of epithelial cells in the pathogenesis of idiopathic pulmonary fibrosis. Cell Signal 66, 109482, doi:10.1016/j.cellsig.2019.109482 (2020).

7 Andrews, D. et al. Unravelling the transcriptional responses of TGF-beta: Smad3 and EZH2 constitute a regulatory switch that controls neuroretinal epithelial cell fate specification. FASEB J 33, 6667–6681, doi:10.1096/fj.201800566RR (2019).

8 Lu, C. et al. Coordination between TGF-beta cellular signaling and epigenetic regulation during epithelial to mesenchymal transition. Epigenetics Chromatin 12, 11, doi:10.1186/s13072-019-0256-y (2019).

9 Skourti-Stathaki, K. et al. R-Loops Enhance Polycomb Repression at a Subset of Developmental Regulator Genes. Mol Cell 73, 930–945 e934, doi:10.1016/j.molcel.2018.12.016 (2019).

10 Tsou, P. S. et al. Inhibition of EZH2 prevents fibrosis and restores normal angiogenesis in scleroderma. Proc Natl Acad Sci U S A 116, 3695–3702, doi:10.1073/pnas.1813006116 (2019).

11 Rubio, K. et al. Inactivation of nuclear histone deacetylases by EP300 disrupts the MiCEE complex in idiopathic pulmonary fibrosis. Nat Commun 10, 2229, doi:10.1038/s41467-019-10066-7 (2019).

12 Grindheim, J. M., Nicetto, D., Donahue, G. & Zaret, K. S. Polycomb Repressive Complex 2 Proteins EZH1 and EZH2 Regulate Timing of Postnatal Hepatocyte Maturation and Fibrosis by Repressing Genes With Euchromatic Promoters in Mice. Gastroenterology 156, 1834–1848, doi:10.1053/j.gastro.2019.01.041 (2019).

13 Yan, J. et al. EZH2 phosphorylation by JAK3 mediates a switch to noncanonical function in natural killer/T-cell lymphoma. Blood 128, 948–958, doi:10.1182/blood-2016-01-690701 (2016).

14 Wan, L. et al. Phosphorylation of EZH2 by AMPK Suppresses PRC2 Methyltransferase Activity and Oncogenic Function. Mol Cell 69, 279–291 e275, doi:10.1016/j.molcel.2017.12.024 (2018).

15 Fernandez, I. E. & Eickelberg, O. The impact of TGF-beta on lung fibrosis: from targeting to biomarkers. Proc Am Thorac Soc 9, 111–116, doi:10.1513/pats.201203-023AW (2012).

16 Zepp, J. A. et al. Distinct Mesenchymal Lineages and Niches Promote Epithelial Self-Renewal and Myofibrogenesis in the Lung. Cell 170, 1134–1148 e1110, doi:10.1016/j.cell.2017.07.034 (2017).

17 Kropski, J. A. & Blackwell, T. S. Endoplasmic reticulum stress in the pathogenesis of fibrotic disease. J Clin Invest 128, 64–73, doi:10.1172/JCI93560 (2018).

18 Subramanian, A. et al. Gene set enrichment analysis: a knowledge-based approach for interpreting genome-wide expression profiles. Proc Natl Acad Sci U S A 102, 15545–15550, doi:10.1073/pnas.0506580102 (2005).

19 Meissner, A. et al. Genome-scale DNA methylation maps of pluripotent and differentiated cells. Nature 454, 766–770, doi:10.1038/nature07107 (2008).

20 Wiederschain, D. et al. Contribution of polycomb homologues Bmi-1 and Mel-18 to medulloblastoma pathogenesis. Mol Cell Biol 27, 4968–4979, doi:10.1128/MCB.02244-06 (2007).

21 Bracken, A. P., Dietrich, N., Pasini, D., Hansen, K. H. & Helin, K. Genome-wide mapping of Polycomb target genes unravels their roles in cell fate transitions. Genes Dev 20, 1123–1136, doi:10.1101/gad.381706 (2006).

22 Pasini, D., Bracken, A. P., Hansen, J. B., Capillo, M. & Helin, K. The polycomb group protein Suz12 is required for embryonic stem cell differentiation. Mol Cell Biol 27, 3769–3779, doi:10.1128/MCB.01432-06 (2007).

23 Kusko, R. L. et al. Integrated Genomics Reveals Convergent Transcriptomic Networks Underlying Chronic Obstructive Pulmonary Disease and Idiopathic Pulmonary Fibrosis. Am J Respir Crit Care Med 194, 948–960, doi:10.1164/rccm.201510-2026OC (2016).

24 Habermann, A. C. et al. Single-cell RNA-sequencing reveals profibrotic roles of distinct epithelial and mesenchymal lineages in pulmonary fibrosis. bioRxiv, 753806, doi:10.1101/753806 (2019).

25 Wei, Y. et al. CDK1-dependent phosphorylation of EZH2 suppresses methylation of H3K27 and promotes osteogenic differentiation of human mesenchymal stem cells. Nat Cell Biol 13, 87–94, doi:10.1038/ncb2139 (2011).

26 Lavarone, E., Barbieri, C. M. & Pasini, D. Dissecting the role of H3K27 acetylation and methylation in PRC2 mediated control of cellular identity. Nat Commun 10, 1679, doi:10.1038/s41467-019-09624-w (2019).

27 Shen, X. et al. EZH1 mediates methylation on histone H3 lysine 27 and complements EZH2 in maintaining stem cell identity and executing pluripotency. Mol Cell 32, 491–502, doi:10.1016/j.molcel.2008.10.016 (2008).

28 Mu, W., Starmer, J., Shibata, Y., Yee, D. & Magnuson, T. EZH1 in germ cells safeguards the function of PRC2 during spermatogenesis. Dev Biol 424, 198–207, doi:10.1016/j.ydbio.2017.02.017 (2017).

29 Aoyama, K. et al. Ezh1 Targets Bivalent Genes to Maintain Self-Renewing Stem Cells in Ezh2-Insufficient Myelodysplastic Syndrome. iScience 9, 161–174, doi:10.1016/j.isci.2018.10.008 (2018).

30 Rana, M. K., Aloisio, F. M., Choi, C. & Barber, D. L. Formin-dependent TGF-beta signaling for epithelial to mesenchymal transition. Mol Biol Cell 29, 1465–1475, doi:10.1091/mbc.E17-05-0325 (2018).

31 Kapoor, P. & Shen, X. Mechanisms of nuclear actin in chromatin-remodeling complexes. Trends Cell Biol 24, 238–246, doi:10.1016/j.tcb.2013.10.007 (2014).

32 Le, H. Q. et al. Mechanical regulation of transcription controls Polycomb-mediated gene silencing during lineage commitment. Nat Cell Biol 18, 864–875, doi:10.1038/ncb3387 (2016).

33 Wei, M. et al. Nuclear actin regulates inducible transcription by enhancing RNA polymerase II clustering. Science Advances 6, eaay6515, doi:10.1126/sciadv.aay6515 (2020).

34 Suh, H. et al. Direct Analysis of Phosphorylation Sites on the Rpb1 C-Terminal Domain of RNA Polymerase II. Mol Cell 61, 297–304, doi:10.1016/j.molcel.2015.12.021 (2016).

35 S. Okuda, N. T., Y. Hasegawa, M. Kawada, Y. Inoue, T. Adachi, Y. Sasai and M. Eiraku. Strain-triggered mechanical feedback in self-organizing optic-cup morphogenesis. Science Advances 4, 12, doi:10.1126/sciadv.aau1354 (2018).

36 Booth, A. J. et al. Acellular normal and fibrotic human lung matrices as a culture system for in vitro investigation. Am J Respir Crit Care Med 186, 866–876, doi:10.1164/rccm.201204-0754OC (2012).

37 Xiao, X. et al. EZH2 enhances the differentiation of fibroblasts into myofibroblasts in idiopathic pulmonary fibrosis. Physiol Rep 4, doi:10.14814/phy2.12915 (2016).

38 Laugesen, A. & Helin, K. Chromatin repressive complexes in stem cells, development, and cancer. Cell Stem Cell 14, 735–751, doi:10.1016/j.stem.2014.05.006 (2014).

39 Mas, G. et al. Promoter bivalency favors an open chromatin architecture in embryonic stem cells. Nat Genet 50, 1452–1462, doi:10.1038/s41588-018-0218-5 (2018).

40 Roberti, A., Valdes, A. F., Torrecillas, R., Fraga, M. F. & Fernandez, A. F. Epigenetics in cancer therapy and nanomedicine. Clin Epigenetics 11, 81, doi:10.1186/s13148-019-0675-4 (2019).

## References

1 Sollner, J. F. et al. An RNA-Seq atlas of gene expression in mouse and rat normal tissues. Sci Data 4, 170185, doi:10.1038/sdata.2017.185 (2017).

2 Ritchie, M. E. et al. limma powers differential expression analyses for RNA-sequencing and microarray studies. Nucleic Acids Res 43, e47, doi:10.1093/nar/gkv007 (2015).

3 Chen, E. Y. et al. Enrichr: interactive and collaborative HTML5 gene list enrichment analysis tool. BMC Bioinformatics 14, doi:10.1186/1471-2105-14-128 (2013).

4 Subramanian, A. et al. Gene set enrichment analysis: a knowledge-based approach for interpreting genome-wide expression profiles. Proc Natl Acad Sci U S A 102, 15545–15550, doi:10.1073/pnas.0506580102 (2005).

5 Mootha VK, L. C., Eriksson KF, Subramanian A, Sihag S, Lehar J, Puigserver P, Carlsson E, Ridderstråle M, Laurila E, Houstis N, Daly MJ, Patterson N, Mesirov JP, Golub TR, Tamayo P, Spiegelman B, Lander ES, Hirschhorn JN, Altshuler D, Groop LC. PGC-1alpha-responsive genes involved in oxidative phosphorylation are coordinately downregulated in human diabetes. Nat Genet. 34, 267–273, doi:10.1038/ng1180 (2003).

6 Cox, J. & Mann, M. MaxQuant enables high peptide identification rates, individualized p.p.b.-range mass accuracies and proteome-wide protein quantification. Nat Biotechnol 26, 1367–1372, doi:10.1038/nbt.1511 (2008).

7 Cox, J. et al. Andromeda: a peptide search engine integrated into the MaxQuant environment. J Proteome Res 10, 1794–1805, doi:10.1021/pr101065j (2011).

8 Le, H. Q. et al. Mechanical regulation of transcription controls Polycomb-mediated gene silencing during lineage commitment. Nat Cell Biol 18, 864–875, doi:10.1038/ncb3387 (2016).

